# Two novel lncRNAs discovered in human mitochondrial DNA using PacBio full-length transcriptome data

**DOI:** 10.1101/079517

**Authors:** Shan Gao, Xiaoxuan Tian, Yu Sun, Zhenfeng Wu, Zhi Cheng, Pengzhi Dong, Qiang Zhao, Bingjun He, Jishou Ruan, Wenjun Bu

## Abstract

In this study, we introduced a general framework to use PacBio full-length transcriptome sequencing for the investigation of the fundamental problems in mitochondrial biology, *e.g.* genome arrangement, heteroplasmy, RNA processing and the regulation of transcription or replication. As a result, we produced the first full-length human mitochondrial transcriptome from the MCF7 cell line based on the PacBio platform and characterized the human mitochondrial transcriptome with more comprehensive and accurate information. The most important finding was two novel lnRNAs hsa-MDL1 and hsa-MDL1AS, which are encoded by the mitochondrial D-loop regions. We propose hsa-MDL1 and hsa-MDL1AS, as the precursors of transcription initiation RNAs (tiRNAs), belong to a novel class of long non-coding RNAs (lnRNAs), which is named as long tiRNAs (ltiRNAs). Based on the mitochondrial RNA processing model, the primary tiRNAs, precursors and mature tiRNAs could be discovered to completely reveal tiRNAs from their origins to functions. The MDL1 and MDL1AS lnRNAs and their regulation mechanisms exist ubiquitously from insects to human.

## Introduction

Animal mitochondrial DNA (mtDNA) is a small, circular and extrachromosomal genome, typically about 16 Kbp in length (16,569 bp for human) and contains almost the same 37 genes: two for rRNAs, 13 for mRNAs and 22 for tRNAs [1], although it’s diversity is being reported. The replication and transcription of human mtDNA is initiated from a noncoding region named Displacement-loop (D-loop). Mitochondrial RNAs are transcribed as primary transcripts, then processed into polycistronic precursors and finally mature RNAs by enzyme cleavage and specific nucleotide modifications. These mature RNAs are pivotal for cellular Adenosine Triphosphate (ATP) production, numerous metabolic regulatory processes, the programming of cell death and other cell functions [2]. However, the exact mechanisms of mitochondrial gene transcription and its regulation are still not well understood [3].

In the year of 1981, the complete human mtDNA sequence was determined and characterized by the encoded genes [4]. With the Next Generation Sequencing (NGS) technologies, Mercer *et al*. tried to demonstrate wide variation in mitochondrial transcript abundance and resolve transcript processing and maturation events [5]. However, the NGS short reads resulted in incompletely assembled mitochondrial transcripts which limited the use of transcriptome data. Based on the Single-molecule Real-time (SMRT) sequencing technology, the PacBio platform can provide longer and even full-length transcripts that originate from observations of single molecules without assembly. Gao *et al*. constructed the first quantitative transcription map of animal mitochondrial genomes [6] by sequencing the full-length insect *(Erthesina fullo Thunberg)* transcriptome [7] on the PacBio platform and built a straightforward and concise methodology to investigate mitochondrial gene transcription. Most of the results in the study of the full-length insect mitochondrial transcriptome were consistent with the previous studies, while new findings included an unexpectedly high level of mitochondrial gene transcription, a high-content of polycistronic transcripts, genome-wide antisense transcripts and a new model of tRNA punctuation *et al*. Using the full-length insect mitochondrial transcriptome data, Gao *et al*. have proved the high-content polycistronic transcripts are mRNA or rRNA precursors and the analysis of these precursors facilitates the investigation of RNA processing, maturation and their mechanisms.

In this study, we used a public PacBio full-length transcriptome dataset to produce the full-length human mitochondrial transcriptome. By the identification and further analysis of full-length transcripts, we were able to characterize the human mitochondrial transcriptome with more comprehensive and accurate information. Our research objectives included: 1) to introduce a general framework for investigating mitochondrial gene transcription using PacBio full-length transcriptome sequencing; 2) to provide accurate annotation of the human mitochondrial reference genome for further studies; 3) to study basic biological similarities and differences between the human and insect quantitative transcription map constructed using their full-length transcriptome data; 4) to reveal molecular mechanisms underlying some fundamental questions, *e.g*. mtDNA heteroplasmy, RNA processing and the regulation of mtDNA transcription or replication.

## Results and Discussion

### The quantitative transcription map of human mitochondrial genome

The full-length transcriptome dataset of the human MCF-7 cell line was chosen to construct the first quantitative transcription map of human mitochondrial genome due to its highest data quality (**Materials and Methods**). This dataset had been acquired by sequencing five groups of cDNA libraries with the size 0~1 Kbp, 1~2 Kbp, 2~3 Kbp, 3~5 Kbp and 5~7 Kbp (**Table 1**). The raw data contained 1,984,154 raw reads with the average size of 16,262 bp. After removing low quality regions and SMRTbell adapters, raw reads were split into 9,192,530 high-quality (Accuracy >= 75%) subreads with the average size of 2,668.85 bp. All the subreads were processed into Circular Consensus Sequencing (CCS) reads to further improve the data quality. Finally, CCS reads were used to produce 744,469 draft transcripts with sequence redundancy. Using adjusted parameters, at least 3.07% (22,853/744,469) of draft transcripts could be continuously mapped to the reconstructed human mitochondrial reference genome (**Supplementary file 1**) to produce the full-length human mitochondrial transcriptome. Since the average length of mitochondrial transcripts is about 1~2 Kbp, the mapping rate was estimated to 7.2% (21,689/301,149) using only data from nine of 28 libraries with the size 0~1 Kbp or 1~2 Kbp. This mapping rate was still significantly lower than the mapping rate 37.65% of the insect *(Erthesina fullo Thunberg)* mitochondrial transcripts [6] (**Figure 1A**). One cause of the difference between these mapping rates could be from the tissue specificity of materials for sequencing. The insect and human total RNA had been extracted from insect whole bodies and human cancer cells, respectively. Other causes could be from the experimental reasons (**Materials and Methods**).

**Figure 1.**
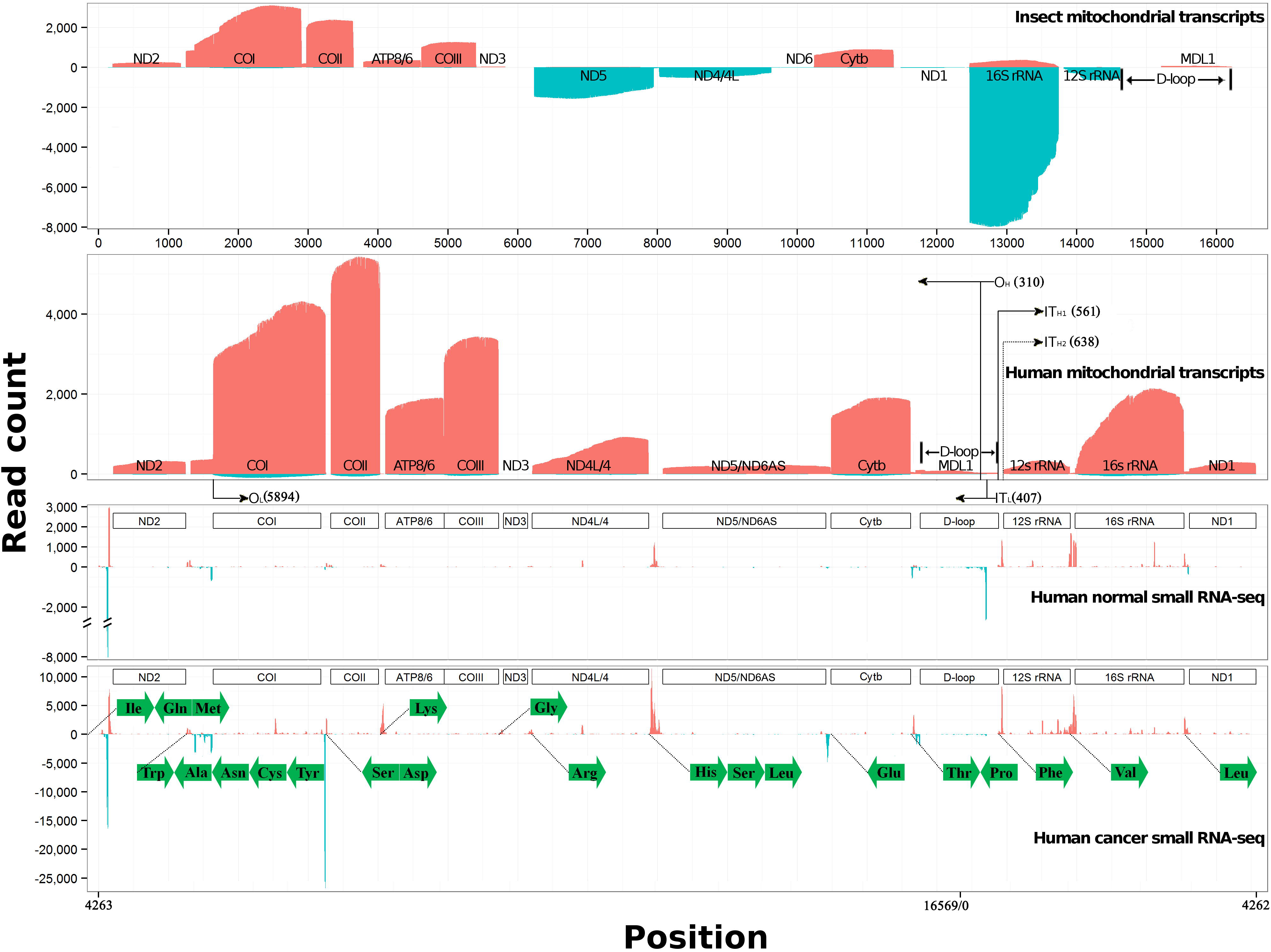
The quantitative transcription map of human mitochondrial genome. A. The quantitative transcription map of insect mitochondrial genome was from our previous study ^9^. Forward alignments of transcripts (in red color) are piled along the positive y-axis. Reverse alignments of transcripts (in blue color) are piled along the negative y-axis. B. The reconstruction of the human mitochondrial genome included two steps. The “N” nucleotide was removed from the complete human mitochondrial genome (Genbank: NC_012920.1) and the resulted sequence shifted 4,263 bp counterclockwise (**Supplementary file 1**). C. Small RNA-seq data of the normal liver tissue (SRA: SRR039612) was aligned to the reconstructed human mitochondrial genome. D. Small RNA-seq data of the Hepatocellular Carcinoma (HCC) tissue (SRA: SRR039625) was aligned to the reconstructed human mitochondrial genome. The tRNA genes (in green color) were represented using their amino acids.

**Table 1.**
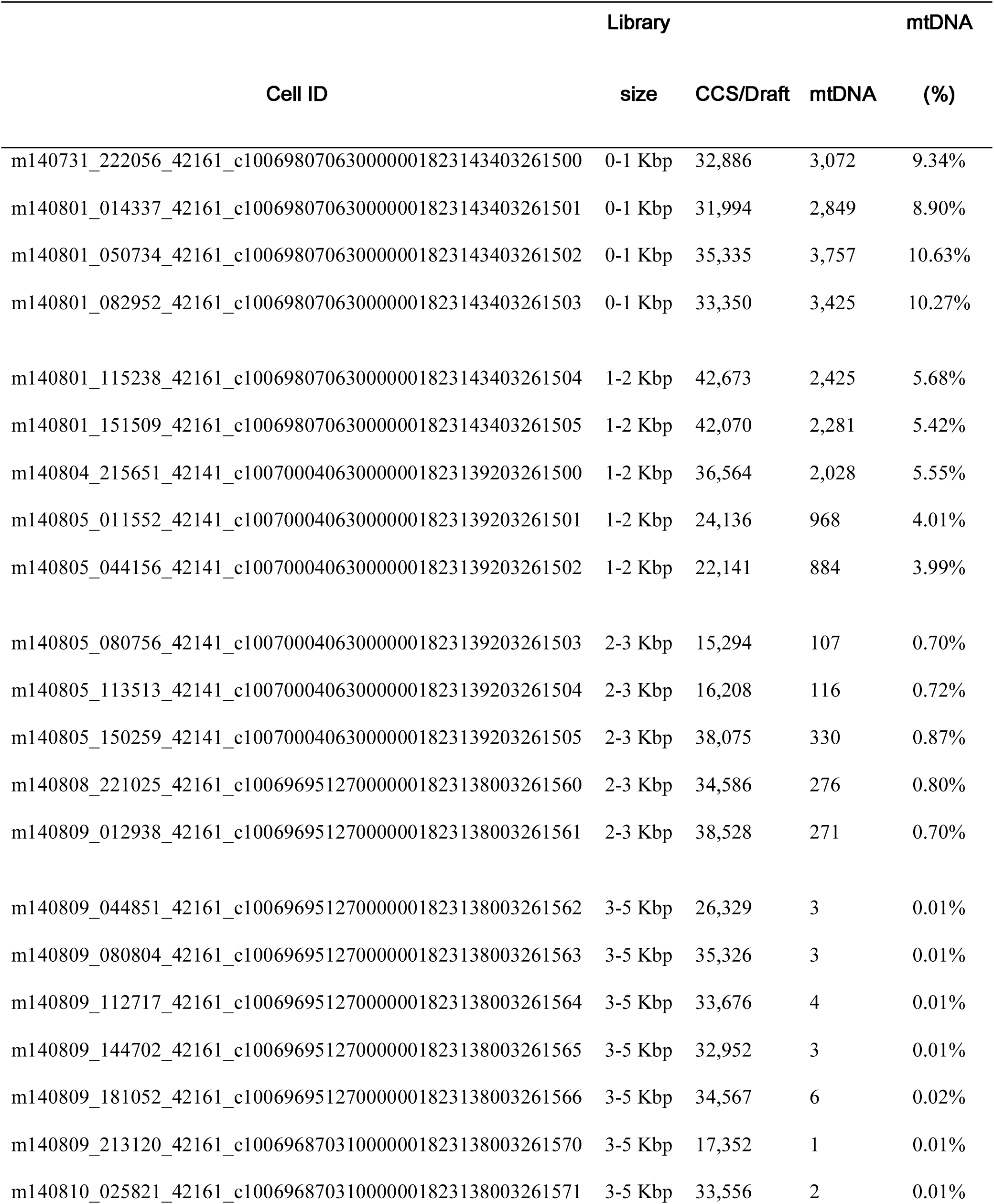
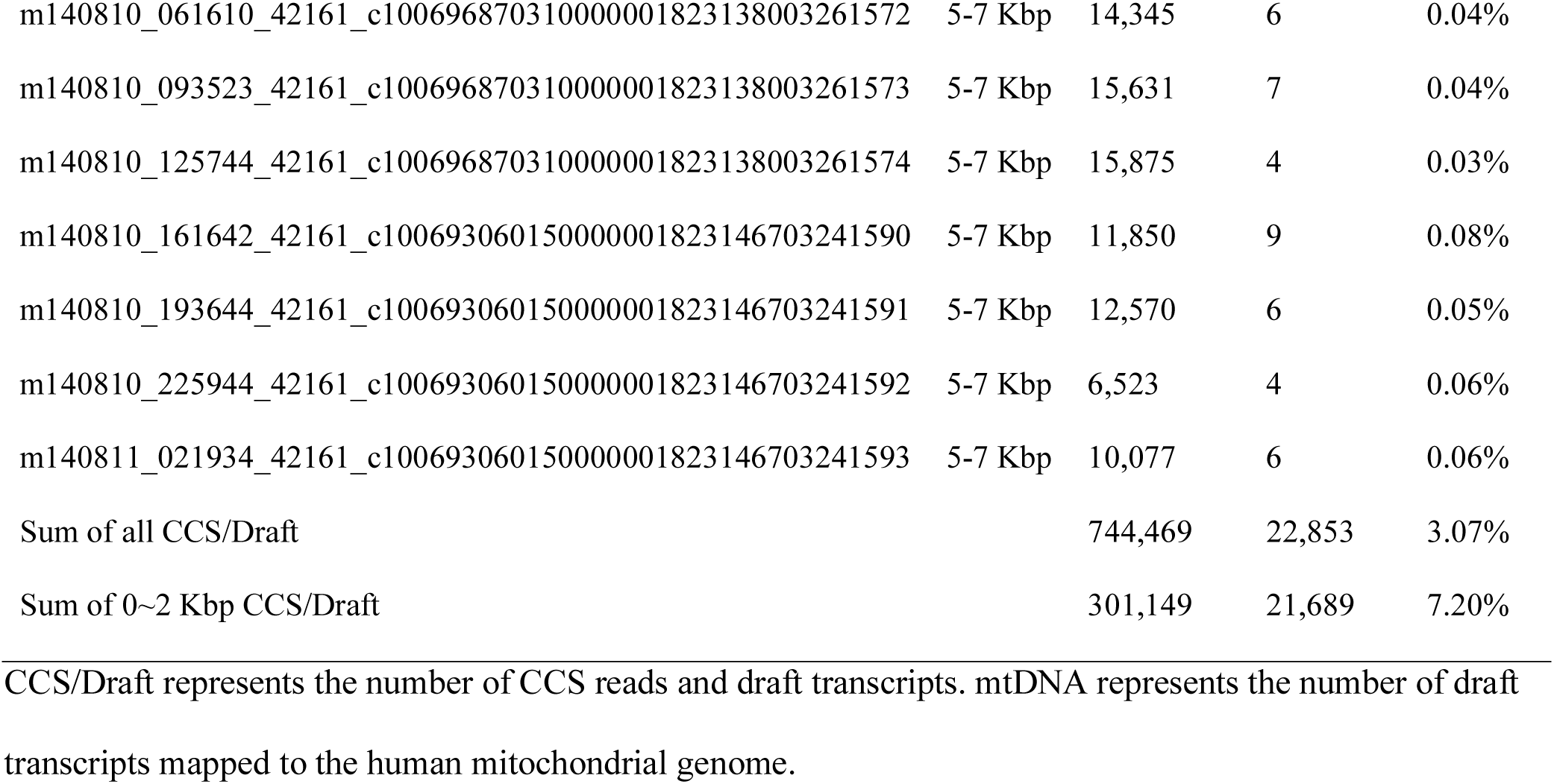
Full-length transcriptome data of human mitochondrial genome

The quantitative transcription map of insect mitochondrial genome (**Figure 1A**) has indicated eight mRNA transcripts (ND2, COI, COII, ATP8/ATP6, COIII, ND3, ND6 and Cytb) are encoded by the major coding strand J(+), also known as the Heavy strand (H-strand) [6], while three mRNA transcripts (ND4L/ND4, ND5 and ND1) and two rRNA transcripts (16S rRNA and 12S rRNA) are encoded by the minor coding strand N(-), also known as the Light strand (L-strand). From **Figure 1B**, it can be seen that all the abundant transcripts (in red color) of the human mtDNA are encoded by J(+), while their antisense transcripts (in blue color) at comparatively low levels are encoded by N (-). It can also be seen that although the ND6 transcript encoded by L-strand is responsible for the protein coding, the antisense ND6 (ND6AS) transcript encoded by H-strand is still more abundant. This finding was different from all the existing human mitochondrial transcription models from the previous studies, in which the ND6AS transcript had been identified as the abundant transcript [8] [4]. Using the NGS data, the ND6 transcript and the antisense ND5 (ND5AS) transcript encoded by L-strand were identified as the abundant transcripts in the year of 2011 [5].

In the full-length human mitochondrial transcriptome, we did not find any full-length transcript of mature ND5 or ND6AS RNAs. This suggested ND5/ND6AS could be one of the fusion genes in the human mitochondrial genome, besides two other fusion genes (ATP8/ATP6 and ND4/ND4L) which had also been found in the insect mitochondrial genome. The human mitochondrial transcripts sorted by transcriptional levels from the highest to the lowest were COII, COI, COIII, 16S rRNA, Cytb, ATP8/ATP6, ND4L/ND4, 12S rRNA, ND2, ND1, ND5/ND6AS and ND3. Although this expression profile had differences from the expression profile of the insect mitochondrial transcriptome due to the tissue specificity and other experimental reasons, the expression profiles of animal mitochondrial mRNA and rRNA genes were still conservative and their transcripts can be roughly classified into high-, medium-and low-level groups according to their transcriptional levels (**Figure 1AB**). In addition, we found eight high-and medium-level mRNA transcripts (COII, COI, COIII, Cytb, ATP8, ATP6, ND4L and ND4) in human mitochondrial genome had the preference to use “ATG” as the start codons, while the other low-level mRNA transcripts (ND2, ND1, ND5 and ND3) had the preference to “ATA” or “ATT”. Since “ATG” has higher efficiency than “ATA” or “ATT” to initial protein synthesis, this suggested the expression of human mitochondrial genes could be regulated coordinately on both of transcriptional and translational levels to ensure the high efficiency.

### Correction of the human mtDNA annotation

The human mitochondrial reference genome (Genbank: NC_012920.1) was annotated with corrections using the identified mature mRNA and rRNA transcripts (**Table 2**). Theese mature mRNA and rRNA transcripts were confirmed by at least 10 full-path CCS reads for their fidelity (**Supplementary file 2**). In general, the identified mature transcripts of the human mtDNA in this study were longer than the predicted ones from the previous study [4]. To determine if the polycistronic ND5/ND6AS transcripts could be further cleavaged into the mature ND5 and ND6AS transcripts, we validated the existence of the mature ND5, ND6AS and ND6 transcripts in different cell lines using qPCR (**Supplementary file 3**). The qPCR results suggested the mature ND5, ND6AS and ND6 transcripts were more likely to contain very short polyA tails [7] than no polyA tails [5]. Since the SMARTScribe reverse transcriptase can catch the transcripts at very low levels, the high depth of Pacbio full-length transcriptome data provides much higher sensitivity than qPCR and RNA-seq in the detection of mature ND5 or ND6AS RNAs. Using a larger Pacbio dataset (**Materials and Methods**), we finally obtained the full-length sequences of the mature ND5, ND6AS and ND6 transcripts (**Table 2**). The ND5 RNA 1 (NC_012920: 12337-14159) and RNA 2 (NC_012920: 1233714163) comprised the most types in the identified mature ND5 sequences (**Figure 2A**), with 11 nt and 15 nt longer than the annotated ND5 RNA (NC_012920: 12337-14148). This 15-nt sequence CCTATTCCCC**<u>C</u>**GAG**<u>C</u>** could contain at least two cleavage sites (underlined). Since the ratio of the sequenced reads supporting the detected ND5 transcripts against the sequenced reads supporting the ND5/ND6AS transcripts was only less than 0.1%, we suspected that the ND5 proteins could be synthesized from both ND5 and ND5/ND6AS transcripts. The mature ND6 transcripts with even lower levels than the ND5 transcripts were identified to have a 27-nt sequence **<u>AGG</u>**TTATGTGAT**<u>TAG</u>**GAG**<u>TAG</u>**GGT**<u>TAG</u>** longer than the annotations (**Figure 2A**). This 27-nt sequence contains three other stop codons “TAG”, besideds the stop codon “AGG” that had been predicted in the previous study [4].

**Figure 2.**
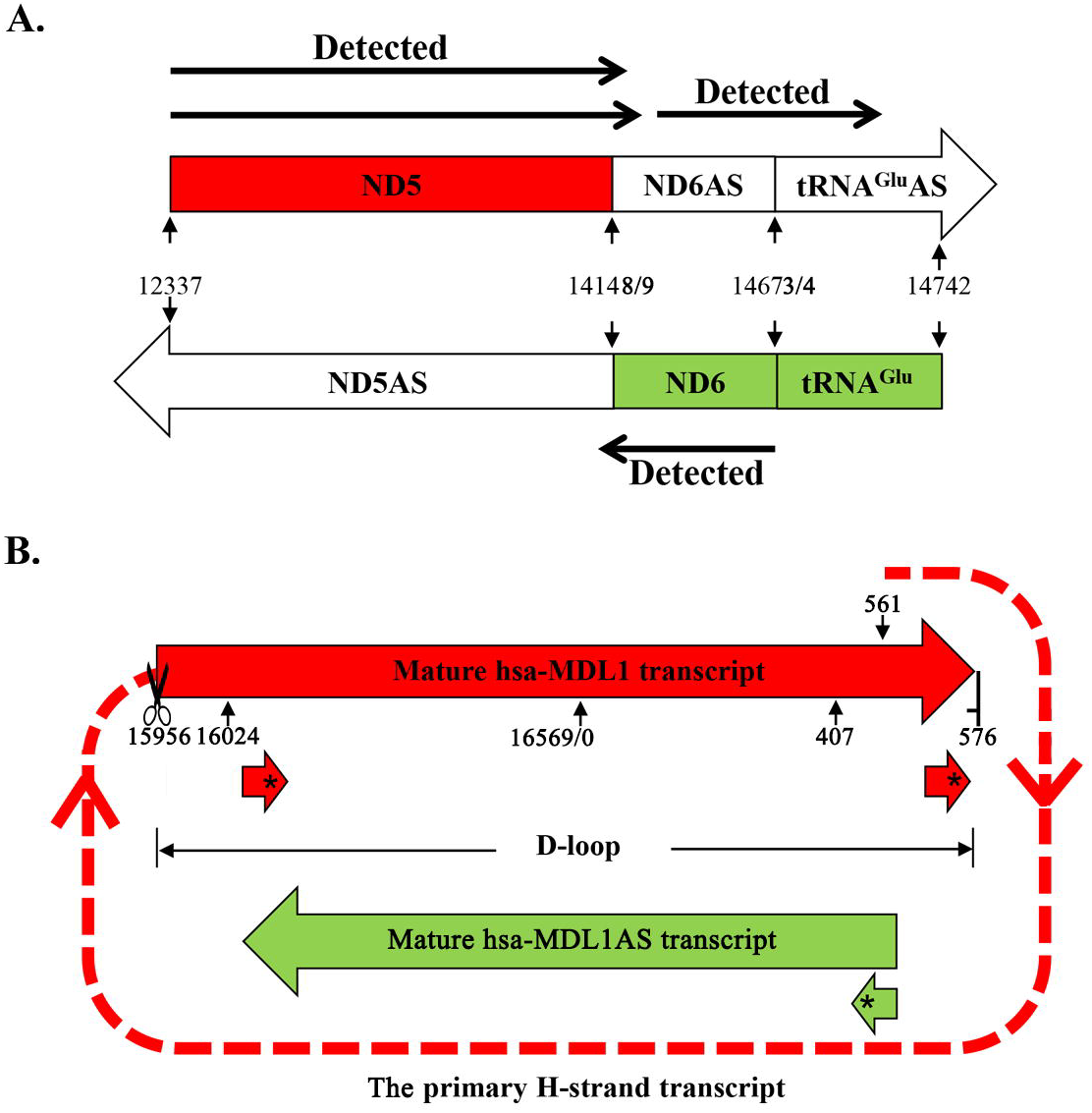
The identified ND5, ND6, MDL1 and MDL1AS transcripts. A. The detected ND5, ND6 and ND6 AS were represented by the arrows with solid lines. The annotated ND5, ND6, ND5AS and ND6AS were marked-by their start and end positions on the human mitochondrial genome (Genbank: NC_012920.1). B. The human D-loop region (NC_012920: 16024-576) encodes the gene hsa-MDL1 (NC_012920: 16024-576) and its antisense gene hsa-MDL1AS (NC_012920: 16024-407). The most abundant aligned small RNAs were marked by *.

**Table 2.** Annotation of the human mitochondrial genome with corrections

| Feature | Strand | Start<sub>old</sub> | End<sub>old</sub> | Start<sub>new</sub> | End<sub>new</sub> | Length |
| --- | --- | --- | --- | --- | --- | --- |
| ND2 | J(+) | 4,470 | 5,511 | 4,470 | 5,511 | 1,042 |
| COI | J(+) | 5,904 | 7,445 | 5,901* | 7,445 | 1,545 |
| COII | J(+) | 7,586 | 8,269 | 7,586 | 8,294* | 709 |
| ATP8/6 | J(+) | 8,366 | 9,207 | 8,365* | 9,207 | 843 |
| COIII | J(+) | 9,207 | 9,990 | 9,207 | 9,990 | 784 |
| ND3 | J(+) | 10,059 | 10,404 | 10,059 | 10,404 | 346 |
| ND4L/4 | J(+) | 10,470 | 12,137 | 10,470 | 12,137 | 1,668 |
| ND5 | J(+) | 12,337 | 14,148 | 12,337 | 14,159* | 1,823 |
|  |  |  |  | 12,337 | 14,163* | 1,827 |
| ND6 | N(-) | 14,149 | 14,673 | 14,122* | 14,673 | 460 |
| Cytb | J(+) | 14,747 | 15,887 | 14,747 | 15,887 | 1,141 |
| D-loop | J(+) | 16,024 | 576 | 16,024 | 576 | 1,122 |
| 12S rRNA | J(+) | 648 | 1,601 | 648 | 1,601 | 954 |
| 16S rRNA | J(+) | 1,671 | 3,229 | 1,671 | 3,228* | 1,558 |
| ND1 | J(+) | 3,307 | 4,262 | 3,305* | 4,262 | 958 |
| ND6AS | J(+) | - | - | 14,164 | 14,746 | 583 |
| MDL1 | J(+) | - | - | 15,956 | 576 | 1,190 |
| MDL1AS | N(-) | - | - | 16,024 | 407 | 953 |
| O<sub>L</sub> | - | - | - | 5,894 | - | - |
| O<sub>H</sub> | - | - | - | 310 | - | - |
| I<sub>L</sub> | - | - | - | 407 | - | - |
| I<sub>H1</sub> | - | - | - | 561 | - | - |
| I<sub>H2</sub> | - | - | - | 638 | - | - |

J(+) and N(-) represents the major and minor coding strand of the mitochondrial genome, respectively. I<sub>H2</sub> is still not well determined. The tRNA genes were represented using their amino acids. \* represented the corrected annotation using the full-length transcriptome data.

The corrected genome annotation also included some important gonomic features that were the origin of H-strand replication (O_H_), the origin of L-strand replication (O_L_), the initiation of the first transcription site on H-strand (IT_H1_), the initiation of the second transcription site on H-strand (IT_H2_) and the initiation of transcription site on L-strand (IT_L_). In this study, O_L_ (NC_012920: 5894), O_H_ (NC_012920: 310), IT_L_ (NC_012920: 407) and IT_H1_ (NC_012920: 561) have been determined, which validated predictions from the previous studies, but IT_H2_ (NC_012920: 638) has not been determined. O_H_ and O_L_ were identified using Single Nucleotide Polymorphisms (SNPs) in the human MCF-7 cell line. Among a total of 17 SNPs (**Supplementary file 3**), 13 single base substitutions were A4769G (ND2), T6776C (COI), A8860G (ATP8), G9966A (COIII), T13260C (ND5), T14319C (ND6), A15326G (Cytb), A15380G (Cytb), C16148T (D-loop), T16519C (D-loop), A263G (D-loop), A750G (12S rRNA) and A1438G (12S rRNA), while four insertion-deletion (InDel) sites included O_L_, O_H_ (D-loop), the position 7527 (tRNA^Asp^) and 2617 (16S rRNA). Among these 17 SNPs, eight were reported in the previous study [5] and O_L_, O_H_ (D-loop), and the position 2617 (16S rRNA) were multi-allelic SNPs, which contributed most to the mtDNA heteroplasmy. Particularly, O_L_ had multiple types of polyC insertion, probably due to L-strand replication errors (**Supplementary file 3**).

Using polycistronic transcripts, IT_L_ and IT_H1_ were identified and confirmed starting at the exact position 407 (D-loop) and 561 (D-loop) on the human mitochondrial genome [8]. The 15-nt promoter element CAAACCCC**<u>A</u>**AAGACA (NC_012920: 553-567) surrounding IT_H1_ (underlined) was also consistent with the finding in the previous study [9]. However, we did not find any full-length transcript to validate H-strand transcription starting at the exact position 638 (tRNA^Phe^), which had been predicted as IT_H2_ in the previous study. According to the dual H-strand transcription model, transcription starts relatively highly frequent at IT_H1_ and lowly frequent at IT_H2_ for the synthesis of two rRNA transcripts and the entire H-strand, respectively [8]. Using a larger Pacbio dataset, we still did not find any sequences to support IT_H2_ and the dual H-strand transcription model.

### Novel genes and mechanisms of gene transcription regulation

In the previous study of the insect mitochondrial transcriptome, we discovered some transcripts aligned forwardly to the D-loop region. The further analysis has indicated these transcripts are encoded by a novel gene Mitochondrial D-loop 1 (MDL1) of *Erthesina fullo* Thunberg (eft) with the length of 1,024 bp. MDL1 (eft-mtDNA: 15210-16233) covers 63.64% (1,024/1,609) of the insect D-loop region (eft-mtDNA: 14625-16233) and contains 11 or 12 repeated units (eft-mtDNA: 15260-16233). This gene with high diversity has two high-frequency mutation sites in each repeated unit (**Supplementary file 3**). In this study, we discovered some polycistronic transcripts (**Supplementary file 2**) aligned forwardly to the human D-loop region. The most abundant one of these transcripts covered the antisense tRNA^Pro^ gene (NC_012920: 1595616023) and 100% of the human D-loop region (NC_012920: 16024-576). Therefore, we named the gene from the insect D-loop region and the gene from the human D-loop region as eft-MDL1 and hsa-MDL1 (NC_012920: 15956-576), respectively. The analysis of all the RNA precursors of hsa-MDL1 supported the mature hsa-MDL1 transcript was cleaved from the long primary H-strand transcript (**Figure 2B**), as the other mRNA and rRNA transcripts were processed. This long primary H-strand transcript started at the position 561 and ended at the position 576 in a 20-nt palindrome sequence AAAGACACCC|CCCAC**<u>A</u>**GTTT (NC_012920: 561-580) overlapping the 15-nt promoter CAAACCCC**<u>A</u>**AAGACA (NC_012920: 553-567) predicted in the previous study. We also discovered a few antisense hsa-MDL1 (hsa-MDL1AS) transcripts (NC_012920: 16024-407). However, we had not found any antisense eft-MDL1 transcript in the previous study. Although the full-length transcripts of MDL1 gene were obtained from both of human and insects, we still validated the hsa-MDL1 transcript using qPCR with Sanger sequencing to rule out the DNA contamination or RNA-DNA chimeric sequences (**Supplementary file 3**) and the hsa-MDL1 and hsa-MDL1AS sequence was submitted to the Genbank database with ID KX859178.

Since the detected transcriptional level of hsa-MDL1AS was much lower than the level of hsa-MDL1, we suspected some hsa-MDL1AS transcripts had been processed into small RNAs for specific biological roles. To validate this hypothesis, we aligned the public small RNA-seq data (**Materials and Methods**) to the human mitochondrial genome and obtained reversely aligned small RNAs enriched in the D-loop region (**Figure 1CD**). Particularly, the 5′ ends of the most abundant small RNA AAAGATAAAATTTGAAAT (NC_012920: 390-407) with the length of 18 nt were precisely aligned to IT_L_ (NC_012920: 407). This sequence was validated to be exist in 931 runs of human small RNA-seq data [10] and named as hsa-tir-MDL1AS-1. The fowardly aligned small RNAs in the human D-loop region were less than 5% of the reversely aligned small RNAs. Most of them were enriched at the 5′ ends or 3′ ends of the hsa-MDL1 gene. These results suggested small RNAs mapped to the D-loop region were the transcription initial RNAs (tiRNAs) [11] and the hsa-MDL1 and hsa-MDL1AS transcripts were their precursors. Since hsa-MDL1 and hsa-MDL1AS are encoded by the transcription initial region and have lengths ~1 Kbp, they are defined as long transcription initial RNAs (ltiRNAs) in this study, which is a novel class of long non-coding RNAs (lnRNAs). We propose that ltiRNAs and tiRNAs consititue a regulation system (**Figure 2B**) to ensure that the expression levels of the mitochondrial genes are able to fluctuate in a large dynamic range. This ltiRNA/tiRNA regulation system have both positive and negative feedback mechanisms to control the expression levels of the mitochondrial genes. Positive feedback can increase the transcription of all the genes to ensure a high productive efficiency. Negative feedback could be used to maintain the expression of mitochondrial genes at normal levels by sense-antisense interactions. In addition, the further analysis of the small RNA-seq data showed the transcriptional levels of the tiRNA hsa-tir-MDL1AS-1 in normal tissues were significantly higher than the levels of hsa-tir-MDL1AS-1 in Hepatocellular Carcinoma (HCC) tissues (**Figure 1CD**). This suggested the ltiRNA/tiRNA system could have a loss of balance control in cancer cells. Since the previous studies demonstrated that the replication of mammalian mtDNA are intimately linked with mitochondrial transcription, the ltiRNA/tiRNA system could also be involved in mtDNA replication. However, the relationship between the mtDNA transcription and replication is still not clear.

### Using RNA precursors to study mitochondrial gene transcription

Our previous study have already proved the high-content of polycistronic transcripts in the full-length transcriptome are mRNA or rRNA precursors and the analysis of these precursors is indispensable for the studies of RNA processing, maturation and their mechanisms. Since the PCR or NGS methods cannot differentiate RNA precursors from their mature transcripts, the analysis of RNA precursors using PacBio data plays an important role in the investigation of mitochondrial gene transcription and its regulation. In our previous and this study [6], we found two main types of RNA precursors ended with mRNAs or rRNAs and proposed the “forward cleavage” model. In this model, the cleavage of tRNAs at their 5′ ends precedes 3′ their ends. Since the current protocol can only capture RNAs with 3′ polyA tails in the full-length transcriptome sequencing, this suggested the 3′ polyadenylation of mature mRNAs or rRNAs could be tightly coupled with the tRNA cleavage at its 5′ ends. In this study, we found a new type of precursors, e.g. ND5/ND6AS/tRNA^Glu^AS (NC_012920: 12337-14746), which ended with antisense tRNAs. This finding supported antisense tRNAs could be recognized by the RNA processing enzymes at their 3′ ends [6], but we did not find this rule worked on MDL1(tRNA^Pro^AS/D-loop). We also found antisense tRNAs could be recognized by the RNA processing enzymes at their 5′ ends by the observation of the full-length COI/tRNA^Ser^AS and COI transcripts. However, our data did not support the ND6AS/tRNA^Glu^AS and ND5/ND6AS/tRNA^Glu^AS transcripts could be processed with further cleavage of tRNA^Glu^AS at the 5′ ends.

In this study, the primary H-strand transcript was precisely determined to start at the position 561 and end at the position 576 (**Figure 2B**). Since the end position was 16 nt after the start position, we were curious if the RNA polymerase has possibilities to read through the termination sites in the D-loop region after the primary H-strand transcript has been completely synthesized. Surperisingly, we found two long sequences to support this “read through” model. The 3853-nt sequence (NC_012920: 14628-1911) spanned the regions of ND6AS, tRNA^Glu^AS, Cytb, tRNA^Thr^, tRNA^Pro^AS, D-loop, tRNA^Phe^, 12S rRNA, tRNA^Val^ and 16S rRNA. Another 1727-nt sequence (NC_012920: 16202-1260) spanned the regions of D-loop, tRNA^Phe^, 12S rRNA. But we did not find any mutation in the termination site to explain this “read through” phenomenon. In addition, we found that the 3853-nt sequence started at the site AA**<u>A</u>**CCCCATTAC (NC_012920: 14626-14637), similar to the 15-nt promoter CAAACCCC<u>A</u>AAGACA (NC_012920: 553567) and ended at the site AAGACCCCCG**<u>A</u>**AACC (NC_012920: 1897-1911), similar to the H-strand termination site AAAGACACCCCCCAC**<u>A</u>**GTTT (NC_012920: 561-580). We susptected that this 3853-nt sequence could be another primary H-strand transcript, which was different from the predicted the primary H-strand transcript spanning the regions of tRNA^Phe^, 12S rRNA, tRNA^Val^ and 16S rRNA.

## Conclusions

In this study, we introduced a general framework to use PacBio full-length transcriptome sequencing for the investigation of the fundamental problems in mitochondrial biology, *e.g*. genome arrangement, heteroplasmy, RNA processing and the regulation of transcription or replication. As a result, we produced the first full-length human mitochondrial transcriptome from the MCF7 cell line based on the PacBio platform and characterized the human mitochondrial transcriptome with more comprehensive and accurate information. The new findings corrected some concepts of mitochondrial gene transcription from the previous studies using qPCR [12] or the Illumina NGS technology [5]. The most important finding was two novel genes MDL1 and MDL1 AS, which were encoded by the mitochondrial D-loop regions. The hsa-MDL1 and hsa-MDL1AS transcript were identified as the precursors of tiRNAs, the most abundant of which was hsa-tir-MDL1AS-1 (AAAGATAAAATTTGAAAT). We propose that hsa-MDL1 and hsa-MDL1AS belong to a novel class of long non-coding RNAs (lncRNAs) and are named as long transcription initial RNAs (ltiRNAs). ltiRNAs and tiRNAs consititue a system to regulate gene expression with new mechanisams.

The phenomenon of tiRNAs was observed but neglected in the NGS data analysis from the beginning. Then, tiRNAs were considered as a new class of small RNAs that were predominantly 18 nt in length and mapped within-60 to +120 nt of Transcription Initiation Sites (TISs) in human, chicken and *Drosophila*. The previous studies also reported tiRNAs could be on the same strand as the TISs and preferentially associated with GC-rich promoters. The characteristics of tiRNAs discovered using the full-length transcriptom data with the small RNA-seq data in this study were consistent with predominantly 18 nt in length but not consistent with the mapped genome range (-60 to +120 nt of TISs), high G+C content and transcription strand preferences discovered in the previous studies. Although tiRNAs were associated to some biological functions or human deseases by several experiments, the theroies and research models of tiRNAs have not be built without the knowledge of their precursors. In this study, we associated tiRNAs to their precursors ltiRNAs and proposed the ltiRNAs/tiRNAs regulation system. Based on the mitochondrial RNA processing model, the primary tiRNAs, precursors and mature tiRNAs could be discovered to completely reveal tiRNAs from their origins to functions. Since the mitochondrial genome originated from a eubacterial (specifically alpha-proteobacterial) ancestor [13], the ltiRNA/tiRNA system exist ubiquitously in eukaryotes and prokaryotes.

## Materials and Methods

### Full-length human transcriptome datasets

MCF-7 is a cancer cell line that was first isolated in 1970 from the breast tissue of a 69-year old Caucasian woman. Full-length transcriptome data of the human MCF-7 cell line were downloaded from the amazon website (http://datasets.pacb.com.s3.amazonaws.com), including an old dataset (released in 2013) using the P4/C2 sequencing reagents and a new dataset (released in 2015) using the P5/C3 sequencing reagents. Since P5/C3 reagents increased the sequencing length, we only used the new dataset for this study. To produce this new dataset, the cDNA libraries of MCF-7 had been prepared using SMARTer PCR cDNA Synthesis Kit (Clontech, USA). Size selection had been implemented using the SageELF device (Sage Science, USA) with every two lanes binned together to create libraries with the size 0-1 Kbp, 1-2 Kbp, 2-3 Kbp, 3-5 Kbp and 5-7 Kbp. These libraries had been sequenced on the Pacific Biosciences RS II sequencer using 28 SMART Cells in the year of 2014. Another larger full-length human transcriptome dataset had been obtained by sequencing of normal brain, heart and liver tissues using 115 SMART Cells. This dataset was downloaded from http://www.pacb.com/blog/data-release-whole-human-transcriptome/ and used to confirm the findings from the MCF-7 dataset.

To conduct transcriptome comparison, we also used the full-length *E. fullo* mitochondrial transcriptome data from our previous study. Since the lengths of mitochondrial rRNA and mRNA transcripts distribute in the range of 1~2 Kbp, the insect mitochondrial cDNAs had been amplified without size selection in the study of the *E. fullo* mitochondrial transcriptome. In this study, the human mitochondrial cDNAs had been amplified with size selection to reduce the bias caused by the SMART Cell loading for sequencing. However, there are many other experimental reasons which could induce bias into the sequencing data. These experimental reasons include the amplification efficiency of cDNAs with different sizes and the data yield of different SMART Cells *et al*. Therefore, cDNA amplification with size selection cannot ensure a more accurate quantification of the full-length transcriptome.

### HCC small RNA-seq dataset

The public small RNA-seq dataset used to identify tiRNAs was downloaded from the NCBI SRA database with ID SRP002272 (**Supplementary file 3**). This dataset included 15 clinical samples including three normal liver tissues, one HBV-infected liver tissue, one severe chronic Hepatitis B liver tissue, two Hepatitis B virus (HBV) positive Hepatocellular Carcinoma (HCC) tissues, one Hepatitis C virus (HCV) positive HCC tissue, one HCC tissue without HBV or HCV and six controls. The data analysis (*e.g*. alignment) was conducted following the procedure used in our previous study [10].

### Data analysis

The human mitochondrial reference genome (Genbank: NC_012920.1) was downloaded from the NCBI Genbank database, which has the same sequence with UCSC hg18. Since the mitochondrial D-loop region had been deemed as a non-transcriptional genomic region before this study, the human mitochondrial reference genome contain an interruptted D-loop region. We had to reconstruct the human mitochondrial reference genome for the convenience of annotation and visualization by two steps. The “N” nucleotide was removed from the reference genome (Genbank: NC_012920.1) and the resulted sequence shifted 4,263 bp counter-clockwise (**Supplementary file 1**).

The IsoSeq™ protocol (Pacific Biosciences, USA) was used to process the sequenced reads to Circular Consensus Sequencing (CCS) reads with parameters (Minimum Full Passes = 1, Minimum Predicted Accuracy = 75), then to produce draft transcripts with parameters (Minimum Sequence Length = 300) by removing the 5′ end cDNA primers, 3′ end cDNA primers and 3′ ployA sequences, which had been identified by the pipeline Fastq_clean [14]. Fastq_clean is a Perl based pipeline to clean DNA-seq [15], RNA-seq [16] and small RNA-seq data [10] with quality control and had included some tools to process Pacbio data in the version 2.0 (https://github.com/gaoshanT/Fastq_clean). The software BWA v0.7.12 was used to align draft transcripts to the reconstructed human mitochondrial genome. Alignment quality control and filtering were performed using in-house Perl programs to remove errors in draft transcripts from the IsoSeq™ protocol. Aligned transcripts with query coverage less than 90% or identities less than 90% were removed to filter out alignments with poor quality. Statistics and plotting were conducted using the software R v2.15.3 with the Bioconductor packages [17]. All the identified transcripts (**Supplementary file 2**) were observed and curated using the software Tablet v1.15.09.01 [18].

## Abbreviations

tiRNA: transcription initiation RNA
ltiRNA: long tiRNA
lncRNA: long non-coding RNA
mtDNA: mitochondrial DNA
D-loop: Displacement-loop
ATP: Adenosine Triphosphate
NGS: Next Generation Sequencing
SMRT: Single-molecule Real-time
CCS: Circular Consensus Sequencing
H-strand: Heavy strand
L-strand: Light strand
LCB: Locally Collinear Block
OH: the origin of H-strand replication
OL: the origin of L-strand replication
IT_H1_: the initiation of the first transcription site on H-strand
IT_H2_: the initiation of the second transcription site on H-strand
ITL: the initiation of transcription site on L-strand
SNP: Single Nucleotide Polymorphism
eft: ***Erthesina fullo*** Thunberg
HCC: Hepatocellular Carcinoma
TIS: Transcription Initiation Site

## Declarations

### Ethics (and consent to participate)

Not applicable

### Consent to publish

Not applicable

### Authors’ contributions

SG and WB conceived and supervised this project. SG, XT and ZW processed the data. YS and BH curated the sequences and prepared all the figures, tables and additional files. SG and QZ drafted the main manuscript. JR and PD revised the manuscript. **All authors have read and approved the manuscript**.

### Availability of data and materials

Full-length transcriptome data of the human MCF-7 cell line can be obtained from the amazon website (http://datasets.pacb.com.s3.amazonaws.com). The public small RNA-seq dataset used to identify tiRNAs can be obtained from the NCBI SRA database with accession number SRP002272.

## Acknowledgements

We appreciate the help equally from the people listed below. They are the graduate student Xiaoran Niu, Hong Chang and Professor Guoqing Liu from College of Life Sciences, Nankai University. Dr. Gloria Sheynkman from Dana-Farber Cancer Institute and Dr. Roberto Lleras from Pacific Biosciences kindly provided the experiment information of the full-length human transcriptome. The data analysis in this study was supported by National Scientific Data Sharing Platform for Population and Health Translational Cancer Medicine Specials.

## References

1 Boore, J.L., Animal mitochondrial genomes. Nucleic Acids Research, 1999. 27(8): p. 1767–1780.

2 Shiota, T., K. Imai, J. Qiu, V.L. Hewitt, K. Tan, H.-H. Shen, N. Sakiyama, Y. Fukasawa, S. Hayat, and M. Kamiya, Molecular architecture of the active mitochondrial protein gate. Science, 2015. 349(6255): p. 1544–1548.

3 Stewart, J.B. and A.T. Beckenbach, Characterization of mature mitochondrial transcripts in Drosophila, and the implications for the tRNA punctuation model in arthropods. Gene, 2009. 445(1): p. 49–57.

4 Anderson, S., A.T. Bankier, B.G. Barrell, M.H.L. de Bruijn, A.R. Coulson, J. Drouin, I.C. Eperon, D.P. Nierlich, B.A. Roe, F. Sanger, et al., Sequence and organization of the human mitochondrial genome. Nature, 1981. 290(5806): p. 457–465.

5 Mercer, T.R., S. Neph, M.E. Dinger, J. Crawford, M.A. Smith, A.-M.J. Shearwood, E. Haugen, C.P. Bracken, O. Rackham, and J.A. Stamatoyannopoulos, The human mitochondrial transcriptome. Cell, 2011. 146(4): p. 645–658.

6 Gao, S., Y. Ren, Y. Sun, Z. Wu, J. Ruan, B. He, T. Zhang, X. Yu, X. Tian, and W. Bu, PacBio Full-length transcriptome profiling of insect mitochondrial gene expression. RNA biology, 2016(just-accepted): p. 00–00.

7 Ren, Y., Z. Jiaqing, Y. Sun, Z. Wu, J. Ruan, B. He, G. Liu, S. Gao, and W. Bu, Full-length transcriptome sequencing on PacBio platform (in Chinese). Chinese Science Bulletin, 2016. 61(11): p. 1250–1254.

8 Montoya, J. and G. Attardi, Identification of Initiation Sites for Heavy-Strand and Light-Strand Transcription in Human Mitochondrial DNA. Proceedings of the National Academy of Sciences of the United States of America, 1982. 79(23): p. 7195–9.

9 Chang, D.D. and D.A. Clayton, Precise identification of individual promoters for transcription of each strand of human mitochondrial DNA. Cell, 1984. 36(3): p. 635–43.

10 Wang, F., Y. Sun, J. Ruan, R. Chen, X. Chen, C. Chen, J.F. Kreuze, Z. Fei, X. Zhu, and S. Gao, Using small RNA deep sequencing to detect human viruses. BioMed Research International, 2016. 2016(2016): p. 2596782.

11 Taft, R.J., E.A. Glazov, N. Cloonan, C. Simons, S. Stephen, G.J. Faulkner, T. Lassmann, A.R. Forrest, S.M. Grimmond, and K. Schroder, Tiny RNAs associated with transcription start sites in animals. Nature genetics, 2009. 41(5): p. 572–578.

12 Sanchez, M.I., T.R. Mercer, S.M. Davies, A.M. Shearwood, K.K. Nygård, T.R. Richman, J.S. Mattick, O. Rackham, and A. Filipovska, RNA processing in human mitochondria. Cell Cycle, 2011. 10(17): p. 2904–16.

13 Gray, M.W., G. Burger, and B.F. Lang, The origin and early evolution of mitochondria (Reviews). Genome Biol2:1018. Genome Biology, 2001. 2(6): p. 181–200.

14 Zhang, M., H. Sun, Z. Fei, F. Zhan, X. Gong, and S. Gao. Fastq_clean: An optimized pipeline to clean the Illumina sequencing data with quality control. in Bioinformatics and Biomedicine (BIBM), 2014 IEEE International Conference on. 2014. IEEE.

15 Wang, Y., Z. Wang, X. Chen, H. Zhang, F. Guo, K. Zhang, H. Feng, W. Gu, C. Wu, and L. Ma, The Complete Genome of Brucella Suis 019 Provides Insights on Cross-Species Infection. Genes, 2016. 7(2): p. 7.

16 Cao, Q., A. Li, J. Chen, Y. Sun, J. Tang, A. Zhang, Z. Zhou, D. Zhao, D. Ma, and S. Gao, Transcriptome Sequencing of the Sweet Potato Progenitor (Ipomoea Trifida (HBK) G. Don.) and Discovery of Drought Tolerance Genes. Tropical Plant Biology, 2016: p. 1–10.

17 Gao, S., J. Ou, and K. Xiao, R language and Bioconductor in bioinformatics applications (Chinese Edition). 2014, Tianjin: Tianjin Science and Technology Translation and Publishing Co.

18 Milne, I., G. Stephen, M. Bayer, P.J. Cock, L. Pritchard, L. Cardle, P.D. Shaw, and D. Marshall, Using Tablet for visual exploration of second-generation sequencing data. Briefings in bioinformatics, 2012: p. bbs012.

